# α-Synuclein strains influence multiple system atrophy via central and peripheral mechanisms

**DOI:** 10.1101/2020.10.16.342089

**Authors:** T. Torre Murazabal, A. Van der Perren, A. Coens, A. Barber Janer, S. Camacho-Garcia, N. Stefanova, R. Melki, V. Baekelandt, W. Peelaerts

**Affiliations:** KU Leuven, Laboratory for Neurobiology and Gene Therapy, Department of Neurosciences, Leuven, Belgium; Institut François Jacob (MIRCen), CEA, and Laboratory of Neurodegenerative Diseases, CNRS, Fontenay-aux-Roses, France; Division of Neurobiology, Department of Neurology, Medical University of Innsbruck, Innsbruck, Austria

**Author notes:** Corresponding authors:* Wouter Peelaerts,; Veerle Baekelandt.

## Abstract

Multiple system atrophy (MSA) is a progressive neurodegenerative disease with prominent autonomic and motor features. Different disease subtypes are distinguished by their predominant parkinsonian or cerebellar signs. The pathognomonic feature of MSA is the presence of α-synuclein (αSyn) protein deposits in glial cells of the central and peripheral nervous system. It is unclear why MSA, that invariably presents with αSyn pathology, is clinically so heterogeneous, why it progresses at varying rates and how neuroinflammation affects disease progression. Recently, it was shown that different strains of αSyn can assemble in unique disease environments but also that a variety of strains might exist in the brain of MSA patients. We therefore investigated if different αSyn strains might influence MSA disease progression. To this aim, we injected two recombinant strains of αSyn in MSA transgenic mice and found that they significantly impact MSA disease progression in a strain-dependent way via oligodendroglial, neurotoxic and immune-related mechanisms. Neurodegeneration and brain atrophy were accompanied by unique microglial and astroglial responses and the recruitment of central and peripheral immune cells. The differential activation of microglial cells correlated with the structural features of αSyn strains both *in vitro* and *in vivo*. By injecting αSyn strains in MSA mice we could more closely mimic a comprehensive MSA phenotype in an experimental setting. This study therefore shows that i) MSA phenotype is governed by both the αSyn strain nature and the host environment and ii) αSyn strains can directly trigger a detrimental immune response related to disease progression in MSA.

## Introduction

Multiple system atrophy (MSA) is a rare neurodegenerative syndrome of unknown etiology. It comprises a group of neurological syndromes, including the Shy-Drager syndrome, olivopontocerebellar atrophy (OPCA) and striatonigral degeneration (SND). Today, OPCA and SND are classified as MSA with predominant cerebellar ataxia (MSA-C) or parkinsonism (MSA-P)^1^. Several years before the appearance of motor symptoms, autonomic features such as urogenital dysfunction or orthostatic hypotension develop during a protracted and prodromal phase^1–4^. These autonomic features are highly variable between patients and underscore the heterogeneity of the disease.

Central to MSA pathology is the accumulation of a-synuclein (αSyn) protein in oligodendrocytes but also in neurons^1, 2^. αSyn is invariably found in insoluble deposits in oligodendrocytes of postmortem MSA brain and identification of glial cytoplasmic inclusions (GCIs) is required for a definite diagnosis of MSA. αSyn-containing inclusions are also found in the brain of people with other synucleinopathies, such as Parkinson’s disease (PD) and dementia with Lewy bodies (DLB), but the conformational properties of αSyn aggregates were shown to be specific for MSA^5^. The structure of MSA fibrils purified from human brain was recently solved via Cryo-EM and was found to be highly organized into β-sheet rich filaments that bundle into a twisted fibril^6^. We and others showed that αSyn assemblies isolated, purified and amplified from MSA brain have different biological activities compared to those isolated from the brain of people with PD or DLB^7–10^. This shows that a structure-function relationship exists within αSyn strains and that they might influence disease phenotypes in different synucleinopathies.

Because of their unique structural properties αSyn aggregates behave as prion strains. MSA strains are highly neurotoxic and amplify *in vivo* via seeded templating of soluble αSyn in oligodendrocytes^5^. Even though it has been shown that MSA strains can efficiently propagate in a permissive environment, it is not known if MSA strains will maintain clonality when propagating in different disease environments where host restriction might inhibit strain propagation or strain neurotoxicity. In the case of human prion diseases, prion strains are not monoclonal^6, 11^. Instead, they comprise a cloud of assemblies often with a dominant strain that is maintained and propagates under host selection^12^. MSA protofilaments were recently shown to display subtle differences in their conformation and were found in varying relative abundance in the cerebellum and putamen of MSA patients, which suggests that a cloud of MSA strains might also exist in MSA brain^13^.

The diversity of αSyn strains in synucleinopathies raises the question if different strains might influence oligodendro-, neuropathological or inflammatory processes, which are central to MSA pathology, and hence disease progression in MSA, but this has never been experimentally tested. We therefore asked if αSyn strains can determine MSA disease outcome. To that aim, we injected two well-characterized but structurally distinct recombinant αSyn strains in transgenic MSA mice that constitutively express αSyn in oligodendrocytes^14^. We found that in an MSA disease environment the two strains, termed ribbons and fibrils, can propagate distinct disease phenotypes. Fibrils caused an aggressive and toxic phenotype with severe myelin loss and neurodegeneration. Ribbons, however, caused a milder neurotoxic phenotype but showed more glial pathology. In addition, αSyn fibrils caused a significant pro-inflammatory response and microglial activation with recruitment of peripheral myeloid and leukocytic cells. Because of these pro-inflammatory features, the introduction of αSyn fibrils into the MSA model resulted in an experimental phenotype that more closely represents the MSA condition.

## Materials and Methods

### Generation and labelling of αSyn assemblies

αSyn fibrils and ribbons were generated as previously described^15^. Oligomeric αSyn was produced by incubating monomeric αSyn 4 °C for 7 days. Oligomeric αSyn was separated from monomeric αSyn by size exclusion chromatography using a Superose6 HR10/30 column (GE Healthcare) equilibrated in phosphate buffered saline (PBS) buffer. αSyn assemblies were labelled by addition of 2 molar equivalent of the aminoreactive fluorescent dye atto-488 (ATTO-Tech GmbH). Labelling was performed following the manufacturer’s recommendations. Unreacted dye was removed by size exclusion chromatography or three cycles of sedimentation and suspension in PBS for oligomers or fibrils and ribbons, respectively. The amount of incorporated dye was assessed by mass spectrometry. The nature of the αSyn assemblies used was routinely assessed using a Jeol 1400 (Jeol Ltd, Peabody, MA) Transmission Electron Microscope after adsorption of the samples onto carbon-coated 200-mesh grids and negative staining with 1% uranyl acetate. The images were acquired with a Gatan Orius CCD camera (Gatan).

### Isolation of primary microglia

Primary microglia were derived from P0-P1 C57BL/6 mouse brain. Briefly, after removal of the meninges, the brains were placed in tubes containing Hanks’ Balanced Salt solution (Sigma-Aldrich). Next, they were incubated with 1% trypsin (Gibco-BRL, Life Technologies) for 10 min at 37°C. Following a mechanical dissociation in DMEM supplemented with DNaseI (Sigma-Aldrich), cells were collected by centrifugation for 10 min at 1200 rpm and re-suspended in DMEM, 10% heat-inactivated FCS and 1% Penicillin-Streptomicin and plated in a 75 cm^2^ culture flask. On days 10-14, to collect microglial cells, the microglia-astrocyte co-cultures were shaken on a rotary shaker at 400 rpm for 3 hours. Microglial cells were plated at a density of 300 000 cells/well in a 12-well plate with coverslips. At DIV 14 cells were treated with the different α-SYN assemblies.

### Recombinant αSyn administration and q-PCR

Cells were treated with the different αSyn assemblies in a concentration of 1 μΜ. Untreated cells and cells incubated with BSA and LPS were used as control. After 24 hours the total RNA was extracted from each well and 1 μg of total RNA of primary microglial cells were reverse-transcribed using the High-Capacity cDNA Archive kit (Applied Biosystems, Carlsbad, USA), according to manufacturer’s instructions. cDNA was used in triplicates as template for q-PCR amplification with specific primers and probes for each microglial marker as described in Table 1. Cycling conditions were 10 minutes at 95°C, followed by 50 cycles of 10 seconds at 95°C and 30 seconds at 55°C. The obtained mRNA levels were normalized to the mRNA levels of HPRT housekeeping gene.

**Table 1.**
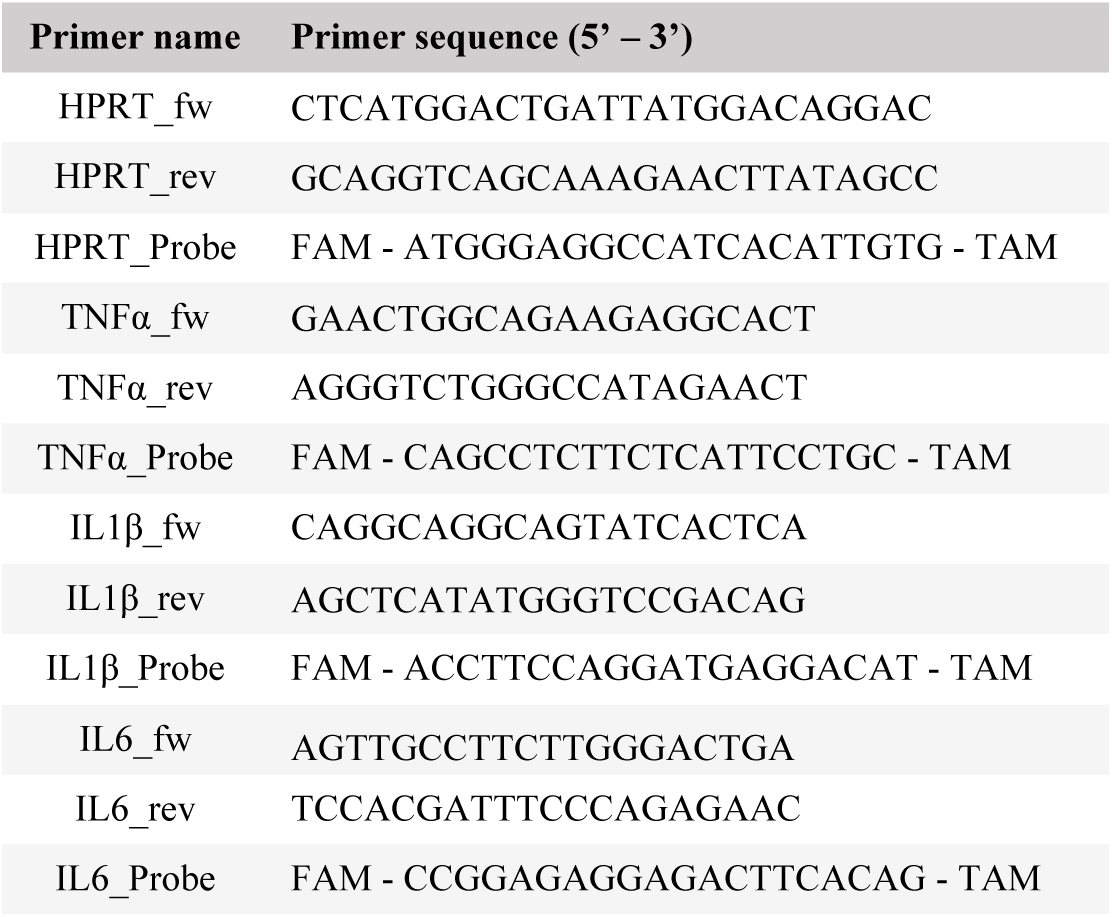
Sequence of primers used for qPCR

### Animals and stereotactic injection

All animal experiments were carried out in accordance with the European Communities Council Directive of November 24, 1986 (86/609/EEC) and approved by the Bioethical Committee of the KU Leuven (Belgium). Approximately 5 month old male and female transgenic (PLP-hαSyn mice^14^) housed under a normal 12-hour light/dark cycle with free access to pelleted food and tap water. All surgical procedures were performed using aseptic techniques and ketamine (70 mg/kg intraperitoneal [i.p.], Ketalar, Pfizer, Belgium) and medetomidine (1 mg/kg, Dormitor, Pfizer) anesthesia. Following anesthesia, the rodents were placed in a stereotactic head frame (Stoelting, IL, USA), a midline incision of the skin was made, and a small hole drilled in the skull at the appropriate location, using bregma as reference. Injections were performed with a 30-gauge needle and a 10μL Hamilton syringe. Animals were injected with 2μL containing 5μg of recombinant protein (BSA, fibrils or ribbons). Stereotactic coordinates used for the dorsal striatum were anteroposterior, +0.5; lateral, −2.0; and dorsoventral, −3.3 calculated from the dura using bregma as reference. The injection rate was 0.25 μL/min and the needle was left in place for an additional 5 minutes before being retracted. Animals were sacrificed after behavioral analysis 9 months post-stereotactic injection.

### Behavioral tests

To examine side bias in spontaneous forelimb use, mice were placed individually inside a glass cylinder (12 cm diameter, 22 cm height). A total of 30 contacts (with fully extended digits executed with both forelimbs) were recorded for each animal. Video-recordings were examined by an observer blinded to the animal’s identity to count the number of touches. The number of impaired forelimb contacts was expressed as a percentage of total forelimb contacts. Non-lesioned control rats should score around 50% in this test. For the pole test, a wooden vertical pole with rope, 1.5 cm of diameter and 50 cm high was used and placed in an open cage. Each mouse was placed with the head up at the top of the pole and the time for turning downwards (T-turn) as well as the total time for climbing down the pole until the mouse reached the floor with the four paws (T-total) was taken in 5 trials. We performed the test 3 times per test and the average of the 3 trials was used for statistical analysis. In the open field (OF), we measured locomotor activity, exploratory and emotional behavior as previously described ^16^. After a 30-minute dark-adaption period, mice were placed in the brightly lit open field area (50 x 50cm^2^). After 1 min of habituation, exploratory behaviour was recorded for 10 min using an automated video tracking system (ANY-maze™ Video Tracking, Stoelting Co. IL, USA).

### Immunocytochemistry

Cells were washed in PBS followed by a permeabilization step in a 0.1% Triton-X100 in PBS solution for 5 min. Next a blocking step of 20 minutes with 10% goat serum in PBS was performed. Cells were incubated with rat anti human αSyn 15G7 primary antibodies (Enzo Life Sciences, 1:500) and rabbit anti-Iba1 (Wako, 1:500) overnight at room temperature. The next day, after 3 washes with PBS cells were incubated with secondary antibody (Alexa Fluor conjugated antibody, 1:500, Molecular probes, Invitrogen) for 1 hour at room temperature. After being rinsed in PBS, coverslips were closed with Mowiol (Calbiochem®, California, US) and DAPI (1:1000). Fluorescent stainings were visualized by confocal microscopy with an LSM 510 unit (Zeiss, Belgium).

### Immunohistochemistry

Mice were anesthetized by intraperitoneal injection of pentobarbital (60 mg/kg, Nembutal, Ceva Sante Animale) and perfused transcardially with saline followed by ice-cold 4 % paraformaldehyde (PFA) in phosphate buffered saline (PBS). Isolation and perfusion were followed by overnight fixation in 4 % PFA. For DAB staining, free-floating sections were pretreated with 3 % hydrogen peroxide (Chem-Lab) in PBS and 10% Methanol for 10 min and incubated overnight with the primary antibody (Table 2) in PBS/T 0.1 with 10 % normal swine/ goat or rabbit serum (Dako). Second, a biotinylated swine anti-rabbit, goat or rabbit (1:300, Dako) was used, followed by incubation with a streptavidin-HRP complex (1:1000, Dako). Immunoreactivity was visualized using DAB (0.4 mg/ml, Sigma-Aldrich) or Vector SG (Vector Laboratories) as a chromogen. After a dehydration series, stained sections were mounted with DPX (Sigma-Aldrich) and visualized with a light microscope (Leica Microsystems).

**Table 2.** Primary antibodies used in this study.

| Antigen | Host | Catalog nr | Brand | Type of detection | Dilution |
| --- | --- | --- | --- | --- | --- |
| CD3 | Rat | ab11086 | abcam | Chromogenic (DAB) | 1:200 |
| CD45 | Rat | sc-1178 | Santa Cruz | Fluorescent | 1:200 |
| CD68 | Rat | MCA1957 | Biorad | Fluorescent | 1:500 |
| GFAP | mouse | 556329 | BD Biosciences | Fluorescent | 1:1000 |
| Iba1 | Rabbit | 019-19741 | Wako | Fluorescent | 1:500 |
| Iba1 | Goat | ab107159 | abcam | Fluorescent | 1:500 |
| MHCII | Rat | 14-532-1-85 | eBioscience | Chromogenic (DAB) | 1:500 |
| NeuN | Mouse | MAB 377 | Millipore | Fluorescent | 1:1000 |
| Olig2 | Rabbit | MAB9610 | Millipore | Fluorescent | 1:1000 |
| P-α-SYN | Rabbit | ab51253 | abcam | Fluorescent | 1:2000 |
| Tmem-119 | Rabbit | ab209064 | abcam | Fluorescent | 1:500 |
| TH | Rabbit | AB152 | Millipore | Chromogenic (Vector SG) | 1:5000 |

For fluorescent double or triple staining, sections were washed in PBS, pre-blocked with 10 % normal horse serum in PBS/T 1 % and incubated overnight with the primary antibodies in PBS/T 1 % with 10 % donkey serum. After washing with PBS, sections were incubated for 2 h with donkey secondary antibodies with different fluorescent tags (1:500, PBS/T 1 %). Next, the sections were washed in PBS and mounted with Mowiol. Fluorescence was detected either with Leica DM6 B automated upright microscope and images were taken using the Leica DFC7000 T camera or by confocal microscopy with a Leica Zeiss LSM 880-Airyscan (Cell and Tissue Imaging Cluster (CIC), Supported by Hercules AKUL/15/37_GOH1816N and FWO G.0929.15 to Pieter Vanden Berghe, KU Leuven). For myelin detection, Fluoromyelin (ThermoFisher, F34651) was used. Sections were incubated in a concentration of fluoromyelin of 1:300 in PBS Triton (1%) for 30 minutes in the dark and washed in PBS.

### Stereological and image quantification

The number of TH-positive cells in the SN was determined by stereological measurements using the Optical fractionator method as described before^17^ (StereoInvestigator; MicroBrightField, Magdeburg, Germany). Every fourth section throughout the SN was analyzed, with a total of 6 sections for each animal. The coefficient of error calculated according to the procedure of Schmitz and Hof (Schmitz and Hof, 2005), varied between 0.05 and 0.10. For the fluorescent triple stereological quantifications, we performed similar stereological measurements, using the same parameters mentioned above but we made use of the software Stereologer®, SRC Biosciences (Stereology Resource Center, Inc.). We quantified both the injected and non-injected SN (internal control). An investigator blinded to the different groups performed all the analyses. For image quantifications we used FIJI software. For fluorescent analyses, we quantified the Mean Fluorescent Intensity (MFI) or the % positive area after adaptive and unbiased thresholding. Threshold was set automatically using either Yen or Triangle threshold. In this case we outline the area of interest and quantify the MFI or percentage of positive area after threshold.

## Results

Transgenic PLP-αSyn mice (from here on referred to as MSA mice) ubiquitously express human αSyn in oligodendrocytes and typically start showing αSyn-rich deposits as early as 2 months^14^. Smaller inclusions, that might represent oligomeric species of αSyn, develop in neurons, astrocytes and microglial cells but never to the same extent as the accumulation of aggresomal, GCI-like structures in oligodendrocytes. Neuroinflammation, characterized by microglial activation, is one of the earliest features that develop in MSA mice together with non-motor features that include loss of neurogenic control of the ^18^ and REM sleep behavior disorder^19^. Autonomic features precede dopaminergic cell loss and motor deficits by several months, which subsequently deteriorate in a progressive manner^14^. During these early and late changes, myelin loss is not typically found, suggesting that in this model demyelination is one of the latest features and that it follows neuroinflammation and neurodegeneration at later stages.

The MSA animal model represents thus several of the clinical features of MSA, and more specifically MSA-P. Many of the phenotypes of these MSA mice are symptoms that occur during both prodromal and late stages of the disease as a result of oligodendro-and neuropathy and the model is therefore well suited to assess whether αSyn strains might affect disease progression in this unique cellular environment. We unilaterally injected 3 μ! of the fibrills or ribbons αSyn strains (5 μg/μl), in the dorsal striatum at 12 weeks of age. As a control group, we injected 3 μ! of BSA at a similar concentration. We allowed the MSA mice to age for 9 months after injection and subjected them to different behavioral tests at 6 and 9 months post injection (p.i.). At 6 months p.i., we did not observe any relative behavioral changes between the different groups injected with the two strains or the control condition (Fig. 1a, b). However, at 9 months p.i., we detected a significant worsening in the pole test in MSA mice injected with fibrils compared to the ribbons and control groups (Fig. 1a, b). Fibril injected MSA mice took significantly longer to turn and descent during the pole test (Fig. 1b). The same conclusion was observed 9 months p.i. using the cylinder test indicating progressive worsening of motor symptoms upon fibril injection (Fig. 1c).

**Figure 1.**
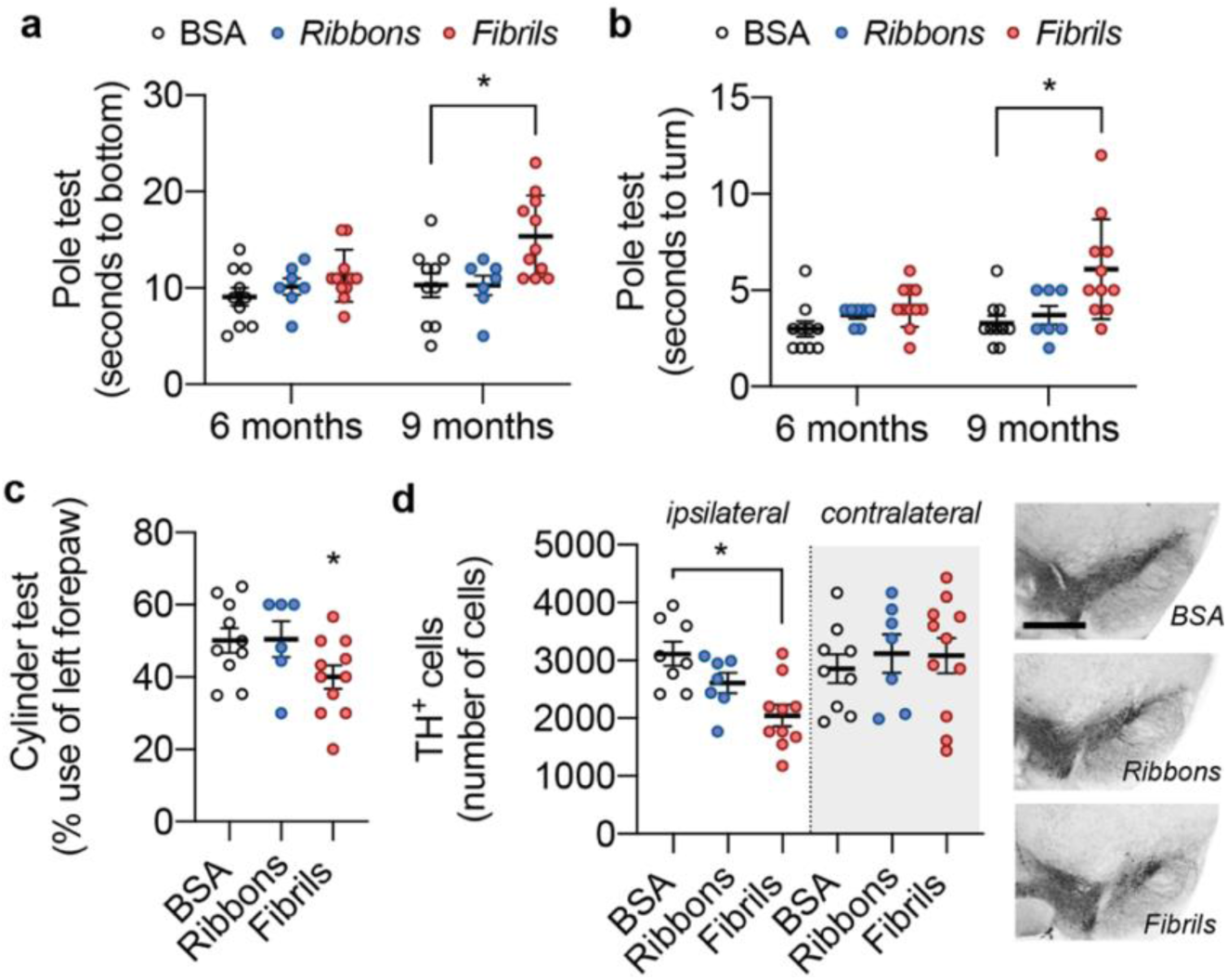
αSyn strains cause distinct behavioral and neuronal pathology in MSA mice. Animals injected with fibrils present progressive motor deficits at 9 months post injection (p.i.). a) Fine motor skills were assessed by the pole test and the time for animals to climb down the pole and b) time to turn (n=7, mean ± SEM, two-way ANOVA with Tukey’s multiple comparison test, *p<0.05, **<0.01). c) Detection of unilateral motor deficits using the cylinder test at 9 p.i. shows a decrease in motor behavior (n=7, mean ± s.e.m., two-way ANOVA Tukey’s Multiple Comparison test, *p<0.05). d) Neurodegeneration is also more prominent in the fibrillar condition with significant dopaminergic cell death compared to MSA mice injected with ribbons or BSA (n=7, mean ± s.e.m., two-way ANOVA with Tukey’s multiple comparison test, *p<0.05.

It has been shown that these motor changes, as assessed by the pole and cylinder tests, reflect neurodegenerative events in the striatonigral pathway. We therefore assessed via stereological quantifications if fibrils could accelerate dopaminergic cell loss in substantia nigra pars compacta (SNpc). Compared to the BSA control condition, we found that fibrils caused significantly tyrosine hydroxylase (TH) cell loss (Fig. 1d). Fibrils worsened TH cell loss by 35%, whereas ribbons caused additional TH cell loss by only 17%. The neurotoxicity of fibrils was further accompanied by a complete loss of fluoromyelin signal throughout the striatum and the corpus callosum (Fig. 2a). The strong loss of myelin was also apparent by severe brain atrophy and ventricle enlargement. Fibrils but not ribbons caused loss of striatal volume of 1.6 mm^3^ (Fig. 2b), corresponding with a ventricular enlargement of 1.4 mm^3^ (Fig. 2c). Given the ubiquitous oligodendroglial expression of αSyn in the transgenic MSA model we could not detect differences in oligodendroglial pSer129-αSyn inclusions between BSA, ribbons or fibrils conditions (Fig. 2d) since every oligodendroglial cell accumulated pSer129-αSyn at this time point. However, when examining pathological accumulation of αSyn, fibrils seeded pSer129-αSyn positive neuritic inclusions in SNpc dopaminergic neurons, whereas ribbons did not (Fig. 2d). Taken together, this shows that fibrils are the most toxic species by seeding MSA pathology resulting in significant worsening of disease progression.

**Figure 2.**
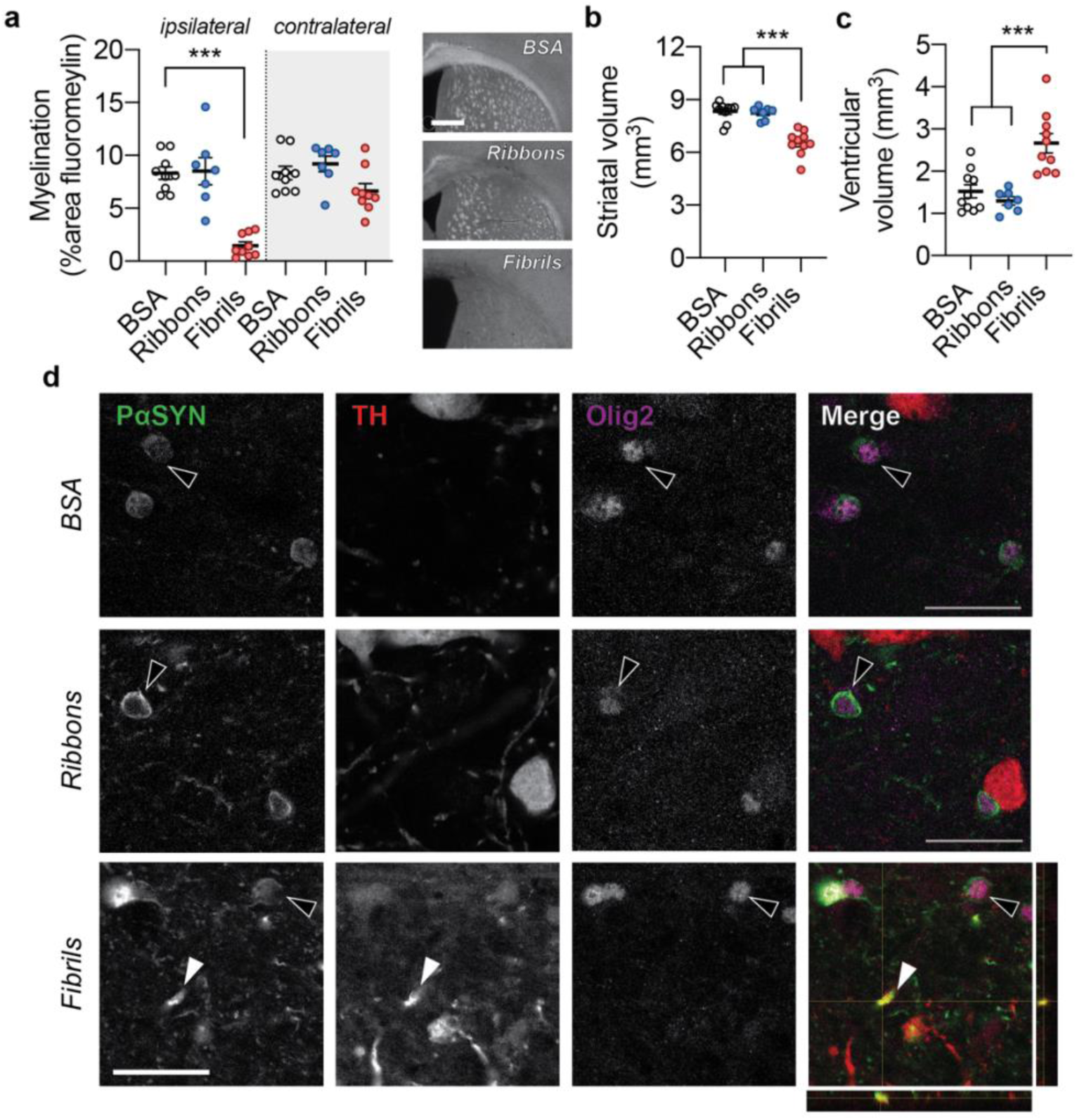
αSyn fibrils causes oligodendrogliopathy with demyelination and brain atrophy. Oligodendrogliopathy was assessed by measuring myelination in the striatum of MSA mice. a) Animals injected with fibrils exhibit severe oligodendroglial pathology whereas animals injected with ribbons or BSA do not (n=7, mean ± s.e.m., two-way ANOVA with Tukey’s multiple comparison test, ***p<0.001). Note the absence of fluoromyelin signal in the corpus collosum and striatum in the fibril condition. Oligodendroglial-and neuropathology in response to the fibrillar strain is further shown by brain atrophy of the b) striatum and c) ventricular enlargement measured (n=7, mean ± s.e.m., two-way ANOVA with Tukey’s multiple comparison test, ***<0.001). d) Representative panel of pSer129-αSyn inclusions in MSA mice injected with BSA, ribbons or fibrils. In all conditions pSer129-αSyn is detected in oliogendrocytes oligodendrocytes (dark arrow). Z-stack from fluorescent staining from the SN of the fibril injected animal shows pSer129-αSyn accumulation in the dopaminergic neurite in addition to GCIs (white arrow) (Scale bar = 25 μm).

To assess if different strains of αSyn can induce immune changes in MSA, we evaluated the effect of the two αSyn strains on microglial function. Inflammation or microglial activation have been shown to be early events in this model of MSA^14^. It is described that different strains of αSyn, including fibrils and ribbons, can elicit a macrophagic response in human monocytes and that this might be strain-dependent^20^. High molecular weight species of αSyn bind to TLR2 and TLR4 receptors, which recognize pathogen-associated patterns, causing glia to become active and produce pro-inflammatory cytokines^21, 22^. In our MSA transgenic mice, TLR4 is upregulated^23^ and microglial cells change from a homeostatic to an active state with the release of pro-inflammatory markers before any detectable neurodegenerative changes have occurred^14^. It is not exactly known what the exact role of macrophagic cells is during the neurodegenerative process, but it is suggested that activated microglial cells might aggravate MSA pathology through the release of pro-inflammatory cytokines and in addition participate in the clearance of the pathogenic αSyn strains^24^. We therefore examined the effects of αSyn strains on the microglial immune response *in vivo* and *in vitro*. Via staining of Iba-1 in combination with the lysosomal marker CD68 for activated microglia we observed strong Iba-1 and CD68 expression in both the fibril and ribbon conditions at 9 months post injection (Fig. 3a-c). Microglial cells were ramified and phagocytic with no clear differences in Iba-1 expression between the two conditions (Fig. 3a-b). In contrast, in the fibril condition we found that microglial cells were significantly more active, as indicated by stronger CD68 staining, whereas this was much less apparent in the ribbons condition (Fig. 3c). Microglial cells were sometimes found to engulf cell debris or pSer129-αSyn inclusions (Fig. 3d) and were furthermore MHCII-positive, especially in the fibrillar condition (Fig. 3e,f) indicating that fibrils are potentially recognized as pathogens and that they can trigger antigen presentation by resident brain macrophages.

**Figure 3.**
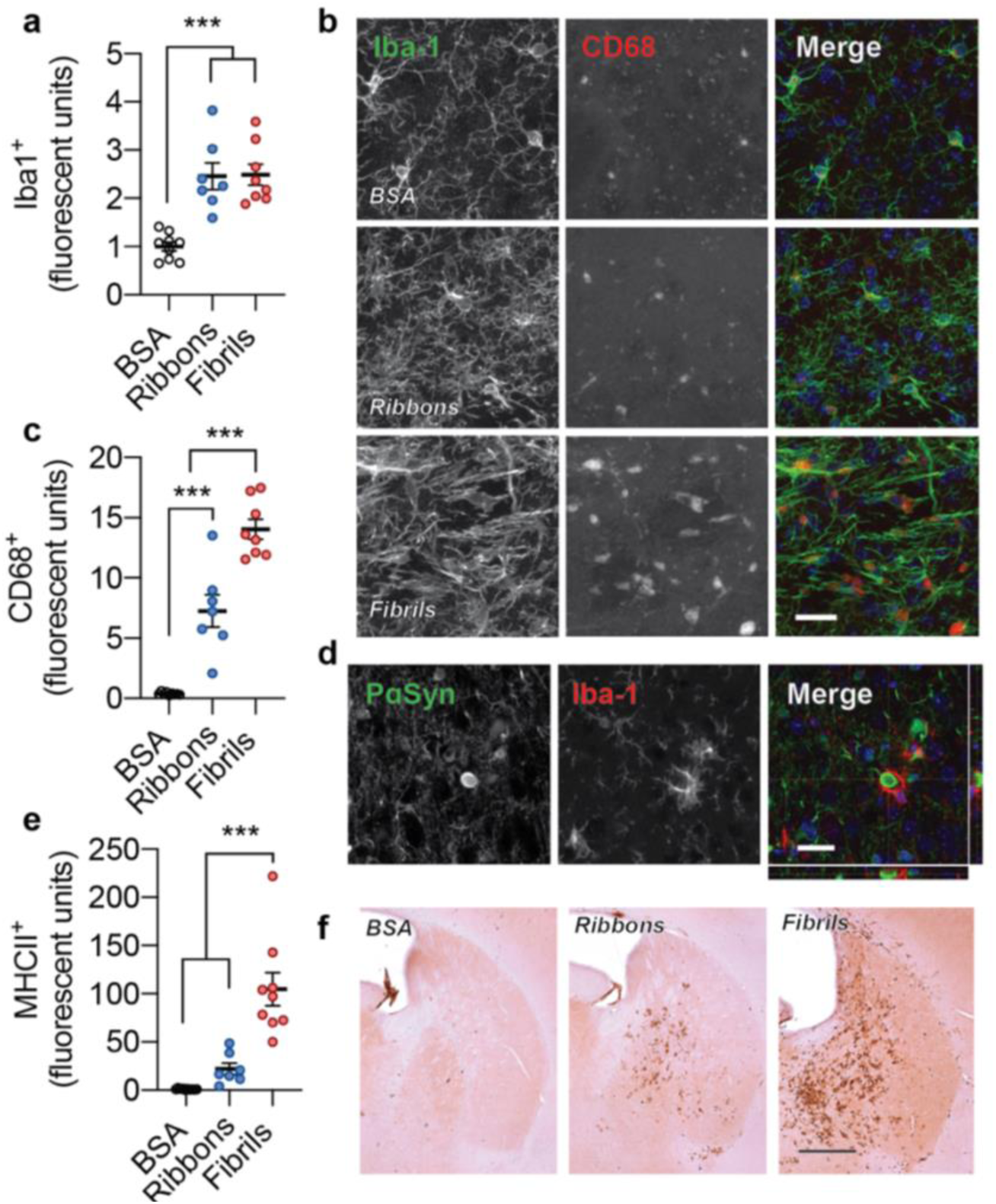
Strain-specific microglial activation in MSA mice. a) αSyn ribbons and fibrils trigger significant microglial activation *in vivo*. Activation was measured via fluorescent positive area after adaptive triangle thresholding (n=7, mean ± s.e.m., one-way ANOVA with Tukey’s multiple comparison test, ***p<0.001). b) Representative images of Iba1 and CD68 staining in the injected striatum in the three experimental conditions (scale bar is 100 μm). c) The microglial lysosomal marker CD68 is significantly upregulated in MSA mice injected with fibrils and ribbons and is indicative of a differential microglial response between the two strain conditions (n=7, mean ± s.e.m., one-way ANOVA Tukey’s multiple comparison test, ***p<0.001). d) Phagocytic Iba1^+^ microglial cells engulfing pSer129-αSyn inclusions (scale bar is 25 μm). e) The microglial response is accompanied by activation and antigen presentation with MHCII expression. f) Fibrils induce strong MHCII expression in the striatum and corpus callosum.

To further define the microglial inflammatory response with respect to the different recombinant αSyn strains, we extended our analysis to primary mouse microglia *in vitro*. We administered different assemblies of αSyn, including monomers, oligomers and the two fibrillar strains to primary murine microglia cultures. Treatment with BSA was used as a negative control. In order to assess uptake of different αSyn assemblies, we performed immunocytochemistry for αSyn and the microglial marker Iba1 at 24 hours after administration. αSyn ribbons and fibrils co-localized with primary microglial cells while monomers and oligomers showed a more diffuse staining pattern (Fig. 4a). Microglial pro-inflammatory response in reaction to different αSyn assemblies was further examined. The expression levels of pro-inflammatory markers TNFα, IL1β and IL6 were strongly upregulated upon administration of αSyn fibrils and to a lesser extent for ribbons. Expression of pro-inflammatory markers was unaffected after treatment with monomeric and oligomeric αSyn (Fig. 4 b-d). This again shows that αSyn strains, and more specifically αSyn fibrils, can act as a direct inflammatory trigger and that the structure of the assembly type is crucial for triggering the immune response.

**Figure 4.**
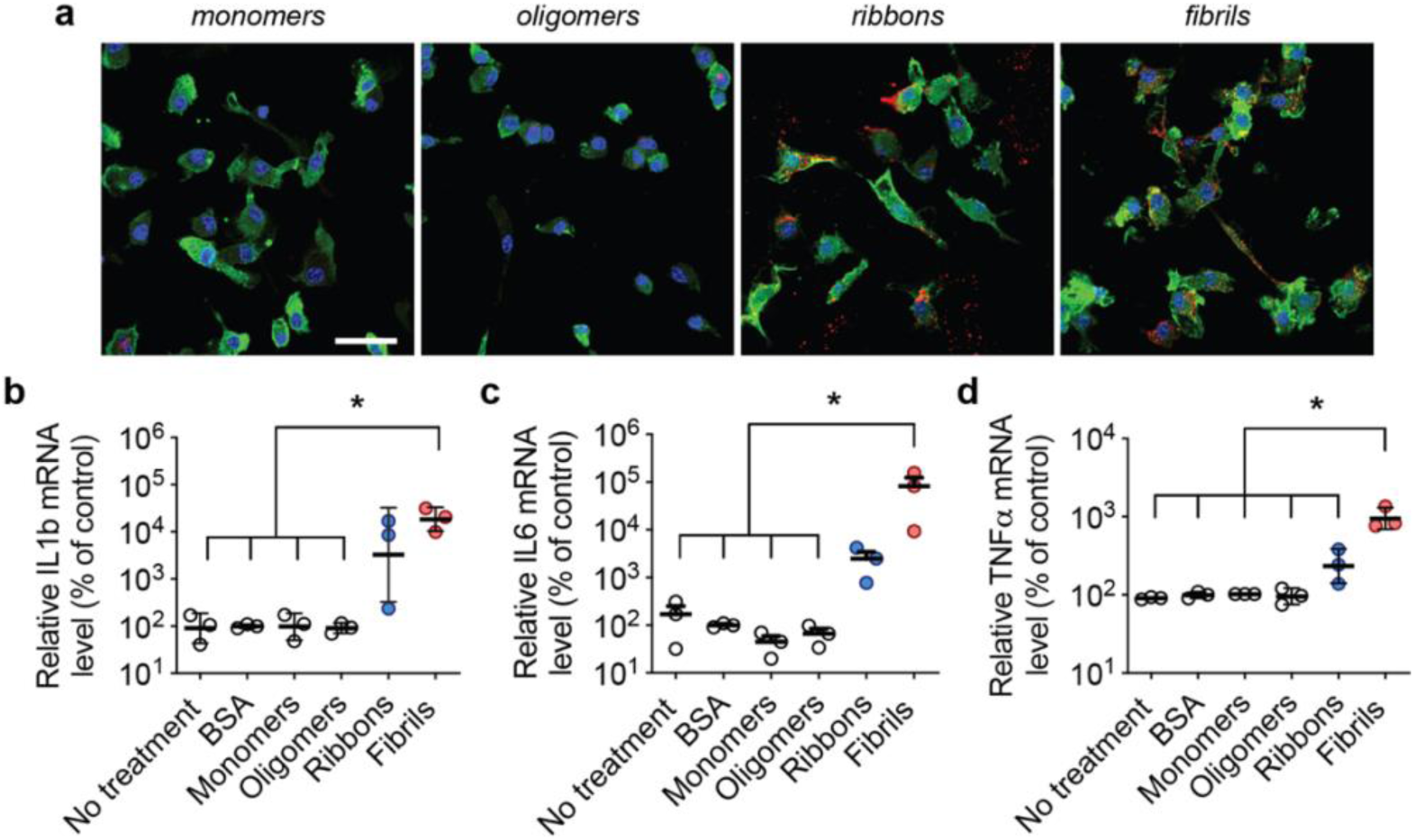
Characterization of the pro-inflammatory response in primary microglia upon treatment with different αSyn assemblies. a) Immunofluorescent staining for human αSyn (red) and Iba1 (green) of primary microglia treated with different αSyn strains for 24 hours. Scale bar represents 50 μm. Quantification of mRNA levels of b) IL1β, c) IL6 and d) TNFα, in murine primary microglia upon administration of monomeric, oligomeric and two different fibrillar αSyn forms (1 μM). Compared to all other tested conditions, αSyn fibrils trigger a more significant pro-inflammatory phenotype. Untreated cells and cells treated with BSA (1 μM) were included as negative controls. Results shown as mean ± SEM, with three unique cell culture experiments at different time points with different assemblies (* p < 0.05, for one-way ANOVA with Tukey’s post-hoc analysis; n=3).

Next to microglial cells, astrocytes take part in the innate immune response and also express different types of TLRs that recognize misfolded αSyn^22, 25^. Extensive astrocytic activation is apparent during post-mortem examination of human MSA brain but is generally absent in the MSA model, suggesting that astrocytic activation might represent an event that occurs at later stages or that the MSA model might require an additional trigger to activate astrocytic response. Astrocytes take up misfolded αSyn via endocytosis as they try to degrade toxic protein via the lysosomal pathway^26, 27^. Unsuccessful clearance of high molecular weight assemblies can sometimes result in protein accumulation and in MSA brain, astrocytic inclusions have been described in various areas including the brain stem and the cerebellum of MSA patients^22–24^. After injecting ribbons and fibrils in the striatum, we found that both strains caused a strong increase of GFAP markers (Fig. 5a-c). Interestingly, even though fibrils were most toxic in MSA mice, they did not cause more astrocytic activation in the striatum compared to ribbons. In addition, only in the ribbon condition, astrocytes were sometimes also positive for pSer129-αSyn (Fig. 5b, d), indicating that astrocytes can take up αSyn strains but also accumulate αSyn as a result of strain exposure or phagocytosis of dyeing cell debris.

**Figure 5.**
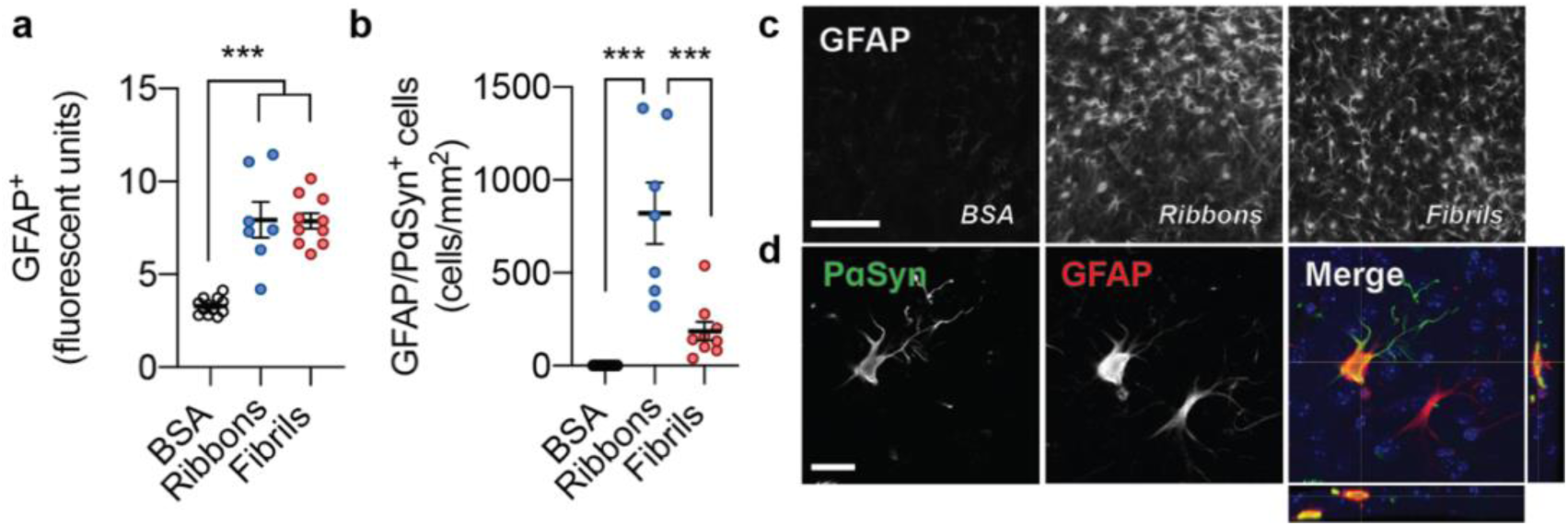
Astrocytic activation and intracellular inclusion formation by αSyn ribbons. a) Striatal injection of ribbons and fibrils in MSA mice cause significant activation of astrocytes (n=7, mean ± s.e.m., two-way ANOVA Tukey’s Multiple Comparison test, ***p<0.001). b) Ribbons induce PαSyn inclusions in astrocytes whereas in the fibril condition this is largely absent (n=7, mean ± s.e.m., two-way ANOVA Tukey’s Multiple Comparison test, ***p<0.001). c) Representative images of GFAP expression of the different experimental and control conditions. d) Colocalization of PαSyn and GFAP shows intracellular glial accumulation of αSyn (scale bar is 25 μm).

Reactive astrocytes and microglia can upregulate cytokine production and release pro-inflammatory mediators that cause neuronal and oligodendroglial damage. Such a pro-inflammatory state could therefore potentially perpetuate the toxicity of αSyn via central but also peripheral immune cells^28^. To confirm whether the Iba-1 cells were resident brain microglial cells, we co-stained for the microgial marker TMEM119 (Iba1^+^/Tmem119^+^), which is absent in peripheral macrophages (Iba1^+^/Tmem119^-^)^29^. Surprisingly, we found significantly higher numbers of non-resident macrophages (Iba1^+^Tmem119^-^) in animals injected with ribbons and fibrils (Fig. 6a, b). To further investigate the potential contribution of peripheral cells, we stained for CD45, a marker highly expressed in peripheral myeloid and leukocytic cells but with low expression in microglial cells. In the atrophied striatum, we found CD45 positive cells throughout the affected area but more abundantly around the lateral ventricles and blood vessels indicating that peripheral immune cells have infiltrated the affected area via these sites (Fig 6c, d). This effect was much less pronounced in the ribbons condition and was absent in the control condition (Fig. 6c). Since we observed that in response to αSyn ribbons and fibrils glial cells became phagocytic active and antigen presenting, next we asked how this might influence the recruitment of T cells. Both in the fibrils and ribbons condition we detected positive staining for CD3 (Fig. 6e, f) suggesting that in conjunction with the presence of MHCII-positive cells T cells might aid microglial cells in the recognition and the clearance of pathogenic species of αSyn. CD3 positive T cells were found throughout the striatum and much more abundant in the case of fibrils compared to ribbons, whereas we did not find any positive cells in the control MSA condition (Fig. 6e).

**Figure 6.**
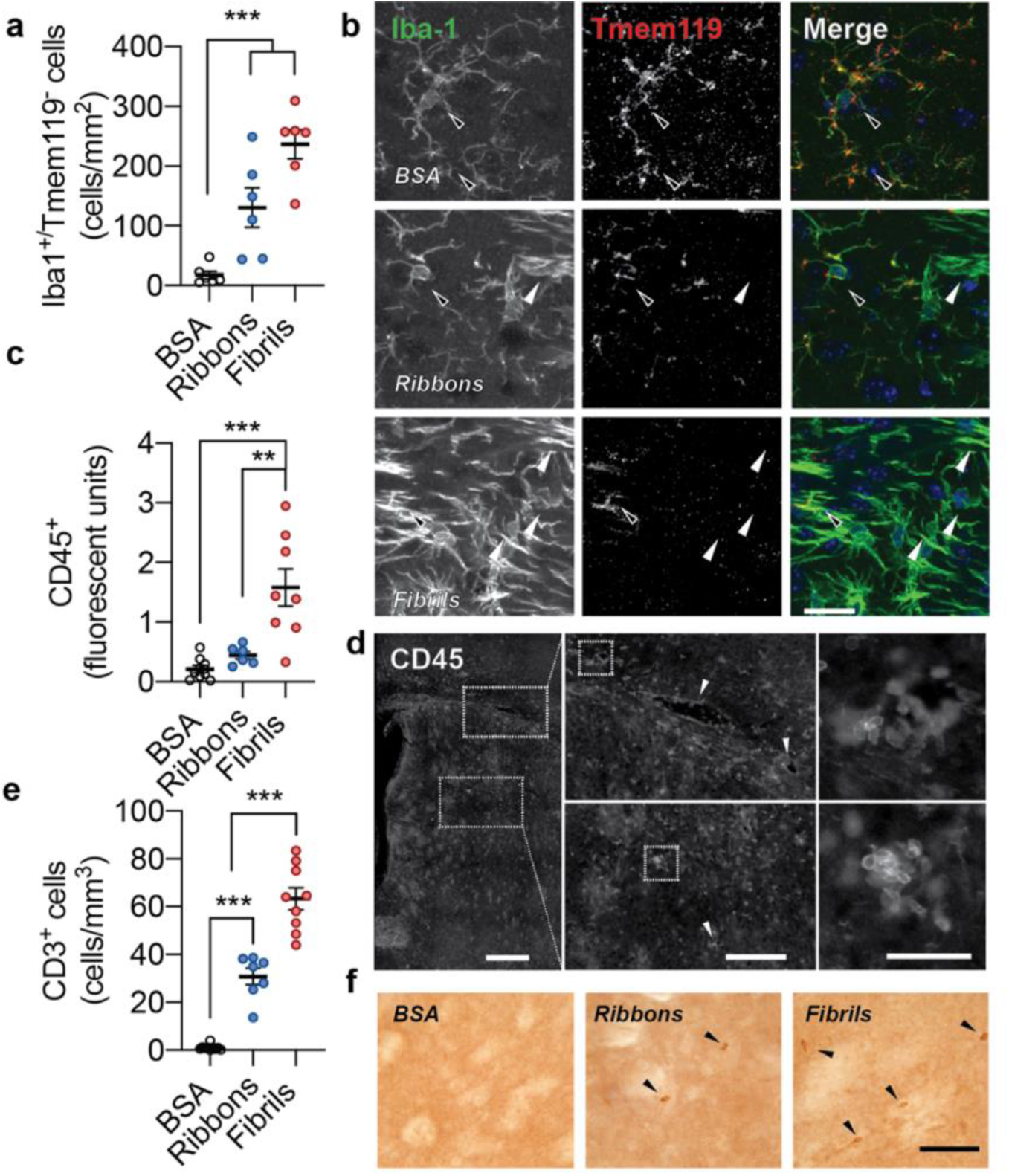
Widespread infiltration of peripheral immune cells in MSA mice after challenge with αSyn fibrils. a) The specific microglial marker Tmem119 allows to distinguish between brain and peripheral macrophages. Co-labeling with Iba1 shows that in the control MSA mice, Tmem119 exclusively colocalizes with Iba-1 (n=7, mean ± s.e.m., one-way ANOVA with Tukey’s multiple comparison test, ***p<0.001). In animals injected with fibrils and ribbons, Tmem119 expression is absent in a subpopulation of Iba-1 cells, suggesting that these macrophages are non-resident immune cells. b) Representative images of Iba1 and Tmem119 in the different conditions showing resident (Tmem119^+^) and peripheral macrophages (Tmem119^-^, white arrows) (scale bar=50μm). c) Peripheral immune cells, detected via the marker CD45, which is highly expressed in all hematopoietic cells, are detected throughout the brain in the fibril condition but are absent in other conditions (n=7, mean ± s.e.m., one-way ANOVA wuth Tukey’s multiple comparison test, **p<0.01, ***p<0.001). d) CD45^+^ cells infiltrate the brain via blood vessels in the corpus callosum and the striatum. Arrows in the middle panel indicate blood vessels and CD45^+^ cells aligned along the vessels. The right panel shows a higher magnification of infiltrating peripheral immune cells. e) Quantification of CD3^+^ T-cells (n=7, Mean ± SEM, one-way ANOVA Tukey’s multiple comparison test, *p<0.05, ****p<0.0001) and f) representative images for infiltrating T-cells.

## Discussion

Recent studies have shown that αSyn strains might play an important role in MSA etiopathogenesis. Through their unique conformation, αSyn strains can cause MSA-like features in cells and *in vivo* ^8^–^30^. Oligodendrocytes offer a unique intracellular environment in which monomeric αSyn forms highly infectious and toxic high molecular weight assemblies^5^. The evidence for a role of strains in MSA disease pathogenesis has mostly been shown via indirect methods but recently it was shown via Cryo-EM αSyn filaments in the brain of MSA patients have defined structural characteristics^13^. Depending on the brain region where αSyn aggregates were isolated from MSA brain it also appeared that small variations existed in the structure of the strain and that the type and the ratio of assemblies varied between patients.

MSA pathogenic assemblies isolated from human brain, either as a crude homogenate or amplified *in vitro,* cause unique neurodegenerative phenotypes upon injections into animals but interestingly, propagation is dependent on the animal model used^5, 8, 31–33^. To date, all inoculation studies performed with MSA strains in WT or transgenic PD mice have unsuccessfully reproduced a robust MSA phenotype. Indeed, we have recently reported that αSyn strains derived from the brain of MSA patients, either as crude homogenates or upon amplification *in vitro*, while inducing the most pronounced disease phenotype in a PD model as compared to PD or DLB strains, do not yield oligodendroglial inclusions^7^. There thus appears to be a discrepancy between the disease strain, the host and the disease phenotype. It has been proposed that synucleinopathies could be triggered by an external event ^34, 35^ but it is not known what such trigger could be in MSA. In the PLP-αSyn mouse model possible external stressors were shown to be oxidative stress/mitochondrial dysfunction^36^ and proteasome disruption^37^. In addition, an incomplete MSA phenotype might arise because of incomplete genetic susceptibility in current animal models, that are often based on PD-specific genes, such as the A53T αSyn mutation^38^, or the absence of a triggering event that facilitates the propagation of MSA strains.

Host genetic susceptibility and exogenous or environmental triggers might thus be more closely related than previously appreciated. In this study we therefore focused on how an MSA disease model can be influenced by exogenous seeds and how different phenotypes might potentially develop. We utilized a well-established transgenic MSA model that mimics parkinsonian features of the disease to assess if distinct strains of αSyn can influence disease phenotype in an MSA environment. By assessing motor behavior, we were able to follow disease progression over multiple months. Motor deficits were reported to occur after 12 months of age in untreated MSA mice and we therefore looked for any progressive features at earlier time points. We found that behavioral motor symptoms appeared at 9 months after striatal fibrils injections, but not in the ribbon condition. In addition, MSA mice injected with fibrils showed severe brain atrophy and cell death whereas this was significantly lower for the ribbons condition. Strikingly, we discovered strong demyelination in multiple disease-associated regions, which previously had only been described in a transgenic MSA model based on overexpression of αSyn by the myelin basic protein promoter^39^.

These results corroborate our previous work where we showed that αSyn fibrils have greater toxicity in vivo compared to ribbons^15, 30^. The reason why fibrils might be more toxic than ribbons can be multifold, but mechanisms of toxicity are governed through the exposed strain surfaces that uniquely interact within their environment^40^. Along those lines, neuroinflammation has been hypothesized to have an important role in MSA. Although its exact role is elusive, microglial activation is considered an early event in MSA etiopathogenesis and persists while the disease progresses. In MSA animals a microglial response is one of the earliest disease events and administration of minocycline, a microglial inhibitor, can ameliorate MSA pathology *in vivo*^23^. Microglial activation accompanies αSyn pathology in human MSA brain, not only in final but also in earlier disease stages^41^ and PET tracers bind with activated microglial cells in putamen and pons of MSA patients^42^. To test if αSyn strains could be involved in these early events as modulators of inflammation, we exposed primary microglial cells to fibrils or ribbons and observed that both strains differentially interact with microglial cells *in vitro*. In contrast to ribbons, fibrils triggered a strong pro-inflammatory response, with the release of Il-ΐβ, Il-6 and TNF-a. Similarly, we found that *in vivo* exposure of fibrils caused severe inflammation *in vivo* with increased phagocytic activity of microglial cells.

Next to microglial cells, astrocytes can take up αSyn fibrils and target it for lysosomal degradation^27^. Astrocytes can assist in the removal of protein aggregates release from neurons and astrocytes are directly coupled via cell junctions to oligodendrocytes^43^. Since a large proportion of binding receptors for fibrils are unique for astrocytes and do not exist in neurons^44, 45^, this raises the possibility that astrocytes might uniquely assist in the removal of defined αSyn assemblies that are released from oligodendroglia. In the MSA mice we found astrocytic inclusions of PαSyn but only in the ribbons group. We could speculate that ribbons might seed inclusions in astrocytes better then fibrils, or alternatively, that fibrils cause rapid cell death of astrocytes resulting in undetectable pathology at the time point studied. Several studies have reported astrocytic inclusions of αSyn in MSA, sometimes in more advanced stages of the disease ^22–24^ Phosphorylated inclusions in astrocytes were found in the ventrolateral part of the spinal cord and brainstem^44^ whereas another study reported astrocytic inclusions in Bergmann glial cells of the cerebellum^45^. In both fibril and ribbon conditions we found activated astrocytic cells, indicated by intense GFAP staining that was absent in the control condition, again showing the importance of strains in αSyn-related pathology.

Altogether, our results demonstrate that αSyn strains can be taken up by immune cells and act as direct triggers of inflammation. Interestingly, neurotoxicity and inflammation in the fibrils condition was also accompanied by strong expression of MHCII receptors on macrophagic cells. This is indicative of an active response towards non self-antigens of abnormal αSyn. Pathogenic assemblies could thus be presented by resident antigen presenting cells that recruit lymphoid cells. Alternatively, αSyn seeds could be secreted in the extracellular space and into the CSF or the lymphatic system where they could activate lymphocytes and elicit a humoral response in a more systemic manner. Recently, it was shown that T-cells can be recruited in a model of experimental MSA^46^. Moreover, they were also detected in post mortem samples of the putamen of MSA patients^46^. In addition, elevated numbers of CD3^+^ and CD4^+^ cells have also been found in the periphery during earlier disease stages^47^. In this study, we now show for the first time and to our best knowledge, that αSyn strains can differentially trigger an adaptive immune response resulting in the recruitment of peripheral lymphocytes. This way, the inflammatory response, triggered by unique αSyn strains in MSA brain, likely contributes to oligodendroglial and neuronal toxicity. This study therefore suggests that inflammation in MSA can be a contributor of pathology in response to αSyn strains. Chronic administration of non-steroidal anti-inflammatory drugs (NSAIDs) can reduce MSA risk ^48^ and cytokine profiling from cerebrospinal fluid and brain tissue from patients showed that pro-inflammatory pathways are upregulated^49–51^. Given that αSyn strains exist in the brain of people with MSA and that strains can specifically cause a detrimental immune response, targeting MSA strains or reducing inflammation in MSA could be a valuable therapeutic strategy.

Additional research will have to investigate what type of immune triggers are important in MSA and if there are activators that are unrelated or downstream from αSyn aggregation. MSA could also be influenced by peripheral infections or inflammation in the gut or the urinary tract which is a frequent feature of MSA^52–54^. Also, there are no clear data on how genetic risk might influence inflammation in MSA. An altered humoral response has been described in PD via association of MHC alleles, which confers increased disease risk^49, 50^. Peptides of post-translationally modified αSyn can elicit an unwanted response from cytotoxic and helper T-cells in PD patients that are recognized by specific MHC alleles^55^. Our work now suggests that the innate and adaptive immune systems are strongly involved in MSA pathogenesis via αSyn strains and that it could potentially influence MSA risk and disease progression. Further investigation will be needed to establish whether an HLA haplotype might exist and influence risk for MSA and whether strains that are specific to MSA could aggravate disease differently, as opposed to PD strains. Collectively, this study demonstrates that the MSA phenotype is influenced by both central and peripheral mechanisms and that the disease phenotype is the consequence of an interaction between the αSyn strain and its cellular environment. Our data also show that via the introduction of disease-relevant strains it is possible to elicit a strong inflammatory response and generate an animal model that more closely mimics the human pathological condition.

## Acknowledgements

We thank L. Bousset for advice in αSyn strains preparation. This work was supported by the FWO Flanders (Projects G.0927.14), the KU Leuven (OT/14/120, C14/18/102), the 2017 and 2020 JiePie MSA-AMS awards to V.B. and W.P., the Flemish Parkinson Liga (VPL) and Fund Sophia managed by the King Baudouin Foundation. W.P. acknowledges a post-doctoral fellowship from Fulbright, IDT Technologies and FWO Flanders. N.S. acknowledges a grant of the Austrian Science Fund (FWF) F4414. R.M. acknowledges support through the JiePie 2019 award. R.M. and A.C. received funding from the EC Joint Programme on Neurodegenerative Diseases (TransPathND, ANR-17-JPCD-0002-02) and the Innovative Medicines Initiative 2 Joint Undertaking under grant agreement No 116060 (IMPRiND). This Joint Undertaking receives support from the European Union’s Horizon 2020 research and innovation programme and EFPIA with support by the Swiss State Secretariat for Education, Research and Innovation (SERI) under contract number 17.00038. The Confocal Images were recorded on a Zeiss LSM 880—Airyscan (Cell and Tissue Imaging Cluster (CIC), Supported by Hercules AKUL/15/37_GOH1816N and FWO G.0929.15 to Pieter Vanden Berghe, KU Leuven).

## Conflict of interest

T.M.T., A.V.P., L.B., A.B.J., S.C., N.S., R.M., V.B. and W.P. report no conflict of interest.

## Notes

### Competing Interest Statement

The authors have declared no competing interest.

## References

1. Fanciulli, A. & Wenning, G. K. Multiple-System Atrophy. N Engl J Med 372, 249–263 (2015).

2. Krismer, F. & Wenning, G. K. Multiple system atrophy: insights into a rare and debilitating movement disorder. Nature Publishing Group 1–12 (2017). doi:10.1038/nrneurol.2017.26

3. Palma, J. A., Norcliffe-Kaufmann, L. & Kaufmann, H. Diagnosis of multiple system atrophy. Autonomic Neuroscience 211, 15–25 (2018).

4. P. G. K. W. et al. The natural history of multiple system atrophy: a prospective European cohort study. The Lancet Neurology 12, 264–274 (2013).

5. Peng, C. et al. Cellular milieu imparts distinct pathological α-synuclein strains in a-synucleinopathies. Nature 1–23 (2018). doi:10.1038/s41586-018-0104-4

6. Schweighauser, M. et al. Structures of α-synuclein filaments from multiple system atrophy. Nature 1–21 (2020). doi:10.1038/s41586-020-2317-6

7. Van der Perren, A. et al. The structural differences between patient-derived α-synuclein strains dictate characteristics of Parkinson’s disease, multiple system atrophy and dementia with Lewy bodies. Acta Neuropathologica 139, 977–1000 (2020).

8. Prusiner, S. B. et al. Evidence for α-synuclein prions causing multiple system atrophy in humans with parkinsonism. Proc. Natl. Acad. Sci. U.S.A. 112, E5308–17 (2015).

9. Shahnawaz, M. et al. Discriminating α-synuclein strains in Parkinson’s disease and multiple system atrophy. Nature 1–23 (2020). doi:10.1038/s41586-020-1984-7

10. Yamasaki, T. R. et al. Parkinson’s disease and multiple system atrophy have distinct α-synuclein seed characteristics. J. Biol. Chem. (2018). doi:10.1074/jbc.RA118.004471

11. Collinge, J. Mammalian prions and their wider relevance in neurodegenerative diseases. Nature 539,217–226 (2016).

12. Li, J., Browning, S., Mahal, S. P., Oelschlegel, A. M. & Weissmann, C. Darwinian Evolution of Prions in Cell Culture. Science 327, 869–872 (2010).

13. Schweighauser, M. et al. Structures of α-synuclein filaments from multiple system atrophy. Nature 1–21 (2020). doi:10.1038/s41586-020-2317-6

14. Refolo, V. et al. Progressive striatonigral degeneration in a transgenic mouse model of multiple system atrophy: translational implications for interventional therapies. Acta Neuropathologica Communications 6, 1–24 (2018).

15. Bousset, L. et al. Structural and functional characterization of two alpha-synuclein strains. Nature Communications 1–13 (2019). doi:10.1038/ncomms3575

16. Callaerts-Vegh, Z. et al. Concomitant deficits in working memory and fear extinction are functionally dissociated from reduced anxiety in metabotropic glutamate receptor 7-deficient mice. J. Neurosci. 26, 6573–6582 (2006).

17. Baekelandt, V. et al. Characterization of Lentiviral Vector-Mediated Gene Transfer in Adult Mouse Brain. 1–13 (2004).

18. Boudes, M. et al. Bladder dysfunction in a transgenic mouse model of multiple system atrophy. Mov Disord. 28, 347–355 (2013).

19. Härtner, L. et al. Distinct Parameters in the EEG of the PLP α-SYN Mouse Model for Multiple System Atrophy Reinforce Face Validity. Front. Behav. Neurosci. 10, 101–13 (2017).

20. Grozdanov, V. et al. Increased Immune Activation by Pathologic α-Synuclein in Parkinson’s Disease. Ann Neurol. 86, 593–606 (2019).

21. Kim, C. et al. Neuron-released oligomeric α-synuclein is an endogenous agonist of TLR2 for paracrine activation of microglia. Nature Communications 4, 382–24 (2013).

22. Fellner, L. et al. Toll-like receptor 4 is required for α-synuclein dependent activation of microglia and astroglia. Glia 61, 349–360 (2012).

23. Stefanova, N. et al. Microglial activation mediates neurodegeneration related to oligodendroglial α-synucleinopathy: Implications for multiple system atrophy. Mov Disord. 22, 2196–2203 (2007).

24. Stefanova, N. et al. Toll-Like Receptor 4 Promotes α-Synuclein Clearance and Survival of Nigral Dopaminergic Neurons. AJPA 179, 954–963 (2011).

25. Rannikko, E. H., Weber, S. S. & Kahle, P. J. Exogenous α-synuclein induces toll-like receptor 4 dependent inflammatory responses in astrocytes. BMC Neuroscience 1–11 (2015). doi:10.1186/s12868-015-0192-0

26. Rostami, J. et al. Human Astrocytes Transfer Aggregated Alpha-Synuclein via Tunneling Nanotubes. J. Neurosci. 37, 11835–11853 (2017).

27. Loria, F. et al. α-Synuclein transfer between neurons and astrocytes indicates that astrocytes play a role in degradation rather than in spreading. Acta Neuropathologica 134, 789–808 (2017).

28. Peralta Ramos, J. M. et al. Peripheral Inflammation Regulates CNS Immune Surveillance Through the Recruitment of Inflammatory Monocytes Upon Systemic α-Synuclein Administration. Front. Immunol. 10, 4784268–6 (2019).

29. Bennett, M. L. et al. New tools for studying microglia in the mouse and human CNS. Proc. Natl. Acad. Sci. U.S.A. 113, E1738–46 (2016).

30. Peelaerts, W. et al. α-Synuclein strains cause distinct synucleinopathies after local and systemic administration. Nature 522, 340–344 (2015).

31. Woerman, A. L. et al. Multiple system atrophy prions retain strain specificity after serial propagation in two different Tg(SNCA*A53T) mouse lines. Acta Neuropathologica 1–18 (2019). doi:10.1007/s00401-019-01959-4

32. Sargent, D. et al. ‘Prion-like’ propagation of the synucleinopathy of M83 transgenic mice depends on the mouse genotype and type of inoculum. J. Neurochem. 143, 126–135 (2017).

33. Bernis, M. E. et al. Prion-like propagation of human brain-derived alpha-synuclein in transgenic mice expressing human wild-type alpha-synuclein. Acta Neuropathologica Communications 1–18 (2015). doi:10.1186/s40478-015-0254-7

34. Wenning, G. K., Stefanova, N., Jellinger, K. A., Poewe, W. & Schlossmacher, M. G. Multiple system atrophy: A primary oligodendrogliopathy. Ann Neurol. 64, 239–246 (2008).

35. Johnson, M. E., Stecher, B., Labrie, V., Brundin, L. & Brundin, P. Triggers, Facilitators, and Aggravators: Redefining Parkinson’s Disease Pathogenesis. Trends in Neurosciences 42, 4–13 (2019).

36. Stefanova, N. et al. Oxidative Stress in Transgenic Mice with Oligodendroglial. American Journal of Pathology 166, 869–876 (2005).

37. Stefanova, N., Kaufmann, W. A., Humpel, C., Poewe, W. & Wenning, G. K. Systemic proteasome inhibition triggers neurodegeneration in a transgenic mouse model expressing human α-synuclein under oligodendrocyte promoter: implications for multiple system atrophy. Acta Neuropathologica 124, 51–65 (2012).

38. Holec, S. A. M. & Woerman, A. L. Evidence of distinct α-synuclein strains underlying disease heterogeneity. Acta Neuropathologica 1–14 (2020). doi:10.1007/s00401-020-02163-5

39. May, V. E. L. et al. α-Synuclein impairs oligodendrocyte progenitor maturation in multiple system atrophy. Neurobiology of Aging 35, 2357–2368 (2014).

40. Melki, R. How the shapes of seeds can influence pathology. Neurobiology of Disease 109, 201–208 (2018).

41. Keisuke, I. et al. Microglial Activation Parallels System Degeneration in Multiple System Atrophy. JNeuropathol Exp Neurol 63, 43 52 (2004).

42. Kübler, D. et al. Widespread microglial activation in multiple system atrophy. Mov Disord. 14, 9–5 (2019).

43. Orthmann-Murphy, J. L., Abrams, C. K. & Scherer, S. S. Gap Junctions Couple Astrocytes and Oligodendrocytes. J Mol Neurosci 35, 101–116 (2008).

44. Nakamura, K. et al. Accumulation of phosphorylated α-synuclein in subpial and periventricular astrocytes in multiple system atrophy of long duration. Neuropathology 36, 157–167 (2015).

45. Piao, Y.-S. et al. α-Synuclein pathology affecting Bergmann glia of the cerebellum in patients with α-synucleinopathies. Acta Neuropathologica 105, 403–409 (2002).

46. Williams, G. P. et al. T cell infiltration in both human multiple system atrophy and a novel mouse model of the disease. Acta Neuropathologica 139, 855–874 (2020).

47. Cao, B. et al. Elevated Percentage of CD3^+^ T-Cells and CD4^+^/CD8^+^ Ratios in Multiple System Atrophy Patients. Front. Neurol. 11, 555–7 (2020).

48. Starhof, C., Hejl, A.-M., Korbo, L., Winge, K. & Friis, S. Risk of Multiple System Atrophy and the Use of Anti-Inflammatory Drugs: A Danish Register-Based Case-Control Study. Neuroepidemiology 54, 58–63 (2020).

49. Rydbirk, R. et al. Cytokine profiling in the prefrontal cortex of Parkinson’s Disease and Multiple System Atrophy patients. Neurobiology of Disease 106, 269–278 (2017).

50. Yamasaki, R. et al. Early strong intrathecal inflammation in cerebellar type multiple system atrophy by cerebrospinal fluid cytokine/chemokine profiles: a case control study. 1–10 (2017). doi:10.1186/s12974-017-0863-0

51. Li, F., Ayaki, T., Maki, T., Sawamoto, N. & Takahashi, R. NLRP3 Inflammasome-Related Proteins Are Upregulated in the Putamen of Patients With Multiple System Atrophy. J Neuropathol Exp Neurol 77, 1050–1065 (2018).

52. Jecmenica-Lukic, M., Poewe, W., Tolosa, E. & Wenning, G. Premotor signs and symptoms of multiple system atrophy. The Lancet Neurology 11, 361–368 (2012).

53. Engen, P. A. et al. The Potential Role of Gut-Derived Inflammation in Multiple System Atrophy. JPD 7, 331–346 (2019).

54. Villumsen, M., Aznar, S., Pakkenberg, B., Jess, T. & Brudek, T. Inflammatory bowel disease increases the risk of Parkinson’s disease: a Danish nationwide cohort study 1977-2014. Gut 68, 18–24 (2019).

55. Sulzer, D. et al. T cells from patients with Parkinson’s disease recognize α-synuclein peptides. Nature Publishing Group 546, 656–661 (2017).

